# The role of social-environmental factors on *Procambarus clarkii* dispersion capabilities

**DOI:** 10.1101/2025.04.09.647993

**Authors:** Daniel Oliveira, Marta C Soares, Pedro Anastácio, Filipe Banha

## Abstract

*Procambarus clarkii* is a prominent example of a successful invasive species, and its ability to disperse is known to be a key characteristic. The ability to disperse dictates the spread and impacts an invasive species has, with various known factors influencing it, such as temperature and circadian rhythms. However, there is still limited information on how crayfish social interactions influence their dispersion, and even already known factors still require further investigation. As such, the goal of this study is to deepen the knowledge of the *Procambarus clarkii* dispersion capabilities. To achieve this, two laboratory experiments were conducted to address the crayfish’ dispersal abilities, namely their speed, under controlled conditions. The first experiment tested the effect of the interaction of two temperatures (16°C and 22°C) and the presence or absence of food in a fully lit or darkened room. The second experiment tested the effects of pairing individuals of the same and opposite sex and of different and/or similar sizes. Three velocity values were obtained: the first considered the crayfish’s final position on the track, the second considered the crayfish’s furthest point on the track, and finally, the third considered the total distance travelled by the crayfish. Results show that when exposed to the higher temperature and in the absence of food, crayfish increased their speed, but size and sex did not seem to have an effect overall. When paired, female crayfish that were behind another crayfish achieved faster speeds on average than males in the same position. We also found that smaller females had faster velocities when paired with bigger females than with bigger males. The observed increase in speed with higher temperatures suggests that, in the face of current global and habitat changes, *P. clarkii*’s rate of spread may accelerate. However, if provided with a reliable food source, crayfish may disperse less. Still, more research is needed to expand our understanding of which factors contribute to *P. clarkii*’s dispersion capabilities, as this may prove important in formulating new plans and strategies to limit its impacts.

## Introduction

Crayfish are among the most widely translocated freshwater species worldwide and their global distribution has been substantially altered by humans, both between and within continents (Taylor 2001, Holdich 2002). This is frequently a problem since non-native crayfish species can affect native species directly by preying on them (Dunn et al. 2009) or by out-competing them for limited resources such as food or shelter (Hill and Lodge 1994, 1999). They can also affect indirectly native communities by altering water turbidity through burrowing activities (Anastácio et al. 2005, Souty-Grosset et al. 2016), reducing aquatic macrophytes and macroinvertebrates (Hill and Lodge 1995, Gherardi and Acquistapace 2007) or by introducing new pathogens (Souty-Grosset et al. 2016).

*Procambarus clarkii* is an omnivore species of the Cambaridae family, native to northeastern Mexico and the southern USA. This polytrophic keystone species can exert multiple pressures on ecosystems and is a prominent example of a successful invasive species with strong individual variations in dispersal distances (Anastácio et al. 2015, Souty-Grosset et al. 2016). It has been introduced in all continents with the exception of Antarctica and Oceania and is currently recorded in 16 European territories (Souty-Grosset et al. 2016). It is considered the most cosmopolitan freshwater crayfish species in the world (Gherardi 2006, Chucholl 2011, Gutiérrez-Yurrita et al. 2017, Lindqvist and Huner 2017). It’s success as an invasive species stems from characteristics such as early maturity, fast growth rate, high offspring number, rapid and adaptable life cycle, flexible feeding strategy, and a wide ability to disperse and to tolerate extreme environmental changes (Hobbs et al. 1989, Alcorlo et al. 2004, Souty-Grosset et al. 2006, 2016, Gherardi 2007, Loureiro et al. 2015, Gutiérrez-Yurrita et al. 2017, de Abreu et al. 2020) in temporary (Gherardi et al. 2002, Alcorlo et al. 2009) or polluted habitats (Gherardi and Barbaresi 2000). It is also a vector of crayfish plague (*Aphanomyces astaci*), which is responsible for large-scale disappearance of native crayfish species (Souty-Grosset et al. 2016), exacerbating its potential as an invasive species.

The longstanding question on what makes any species a successful invader has been a conundrum of increasing interest to ecologists (Kolar and Lodge 2001, Forsyth et al. 2004). At this point it is known that the ability to disperse may be a key factor in determining the invasion success as well as its impact and rate of spread (Ehrlich 1989, Lodge 1993). Many studies have approached this topic, for ex.: (Aquiloni et al. 2005, Marques et al. 2015, Ramalho and Anastácio 2015, Carvalho et al. 2022, Galib et al. 2022), increasing knowledge about relevant factors for *P. clarkii* dispersion, such as temperature, circadian rhythms and population densities, among others. Gherardi et al. (2000) has shown that *P. clarkii* increases motor activity in water ditches with a rise in water temperature between 5°C to 25°C, while Anastácio et al. (1999) reported increased motor activity between 20°C to 28°C. *P. clarkii was* shown to be more active during night periods (De Miguel and Aréchiga 1994, Correia 1998, Miranda-Anaya 2004, Ramaliro 2012). According to De Miguel and Aréchiga, (1994) *P. clarkii* has outbursts of movement that can last for hours when food is presented to them. Additionally, *P. clarkii* has been shown to disperse overland (Penn 1943, Ramalho and Anastácio 2015). More recently Ramalho and Anastácio (2015) analysed the effect of environmental variables (humidity, temperature, illuminance, vegetation, and slope) on the movement direction and observed that *P. clarkii* tends to move downhill and toward cooler areas when out of water.

Though the effect of some environmental factors on the locomotion of *P. clarkii* is known, there are other factors besides these, such as their social dynamics, that could also influence crayfish dispersion. Size, sex or hierarchical positions may influence which individuals are more likely to spread into new territories. Past studies, most of which focus on dominant/subordinate interactions, suggest that crayfish social interactions may affect their locomotion (ex.: Hayes 1975; Huber and Delago 1998; Issa et al. 1999). For example, Issa et al. (1999) have shown that larger crayfish tend to approach others more often and fight for longer, while smaller crayfish tend to move away or quickly retreat, fleeing from fights or avoiding them entirely. Ranta and Lindström (1992) have shown that larger crayfish are able to evict smaller ones from their burrows. However, studies focussing on how these interactions affect the dispersion are still lacking. All these studies have given us insights on *P. clarkii* dispersion and locomotion, but there is still much to uncover. The role of crayfish social interactions on *P. clarkii* dispersion and locomotion is still unclear and even the effects of better understood factors, such as temperature and circadian rhythms, may differ between populations.

Here we aimed to deepen the understanding of which factors influence *Procambarus clarkii* dispersion capabilities. To achieve this goal, two laboratory experimental designs were used. The first aimed to test three environmental variables, namely temperature (16°C and 22°C), light levels (illuminated and dark) and food (presence or absence of food items), while also testing for the influence of size, sex and physical condition. The second experiment tested if the pairing between individuals of the same and opposite sex of different and similar sizes may lead to differences in their dispersion capabilities.

## Methods

### Crayfish capture

*Procambarus clarkii* individuals were captured in the Marateca stream next to rice fields near the town of Cabrela, Portugal (38.58561 N, 8.528580 W). These were captured on three separate occasions: 21st of November 2022, 12th of May 2023 and 14th of July 2023, with the aid of dip nets for juveniles below 25 mm and Swedish-type crayfish traps baited with sardines for adults (Dorn et al. 2005).

Crayfish were maintained in an acclimatised room at 20°C, inside tanks filled with a depth of ∼6 cm of tap water and kept oxygenated using an aquarium pump. Several PVC tube sections were scattered on the bottom of the tanks to serve as shelter and crayfish were fed with carrots. Approximately every six days the water was changed, and the tanks were cleaned.

Crayfish weight and cephalothorax length were measured to help select individuals and form groups with similar size and weight distributions among them. Crayfish condition was calculated using Fulton’s condition index with the formula K = W/L^3^, where W is wet weight (g) and L is Cephalothorax length (mm) (Ricker 1975). The selected crayfish were then stored individually inside tagged containers (20×20×12 cm) for a minimum of 5 days before experimentation. These containers were filed with dechlorinated tap water up to the top of the individual’s cephalothorax and crayfish were fed carrots prior to the experiment. The containers were cleaned, and the water and uneaten food were replaced with fresh water and food every two days.

### General experimental design

The general set-up consisted of a two-meter-long metal gutter filled with a depth of ∼6 cm tap water that was replaced a day before every experiment (Figure 1). A substrate consisting of medium coarse sand with approximately 1cm of dept was used to cover the track’s bottom. A removable translucent plastic plate separated the initial 10 cm of the track from the rest of it. The track and plastic plate were washed clean the day before each experiment day and the water and sand substrate were replaced. Crayfish went through 3 days of fasting and were acclimated to each temperature for 24 hours before the day of the experiment. On the day of each experiment a few drops of water from each individual containers where the crayfish were kept were mixed with the water inside the track 10 minutes before the first replicate started. This was done to scent the water pre-emptively and prevent the possibility that crayfish might behave differently because of the scent from the previously tested individuals. The water pH, percentage and ppm of dissolved oxygen and temperature were noted before and after the experiments. For each replicate, an individual was placed on the first 10 cm of the track and for 60 seconds it was kept from roaming by the translucent plastic plate, so to acclimatize to the new environment. Following 60 seconds, the plastic plate was removed to allow the individual to roam the gutter freely for a total of 120 seconds or until they reached the opposite end of the track, after which their position or the time they took to reach the end of the gutter was noted. The dispersal capability of individuals was assessed by measuring their velocities. Three velocity values were calculated based on the crayfish final position on the track (final point velocity), crayfish furthest point on the track (furthest point velocity) and crayfish total distance travelled (real velocity) (Figure 1).

**Figure 1:**
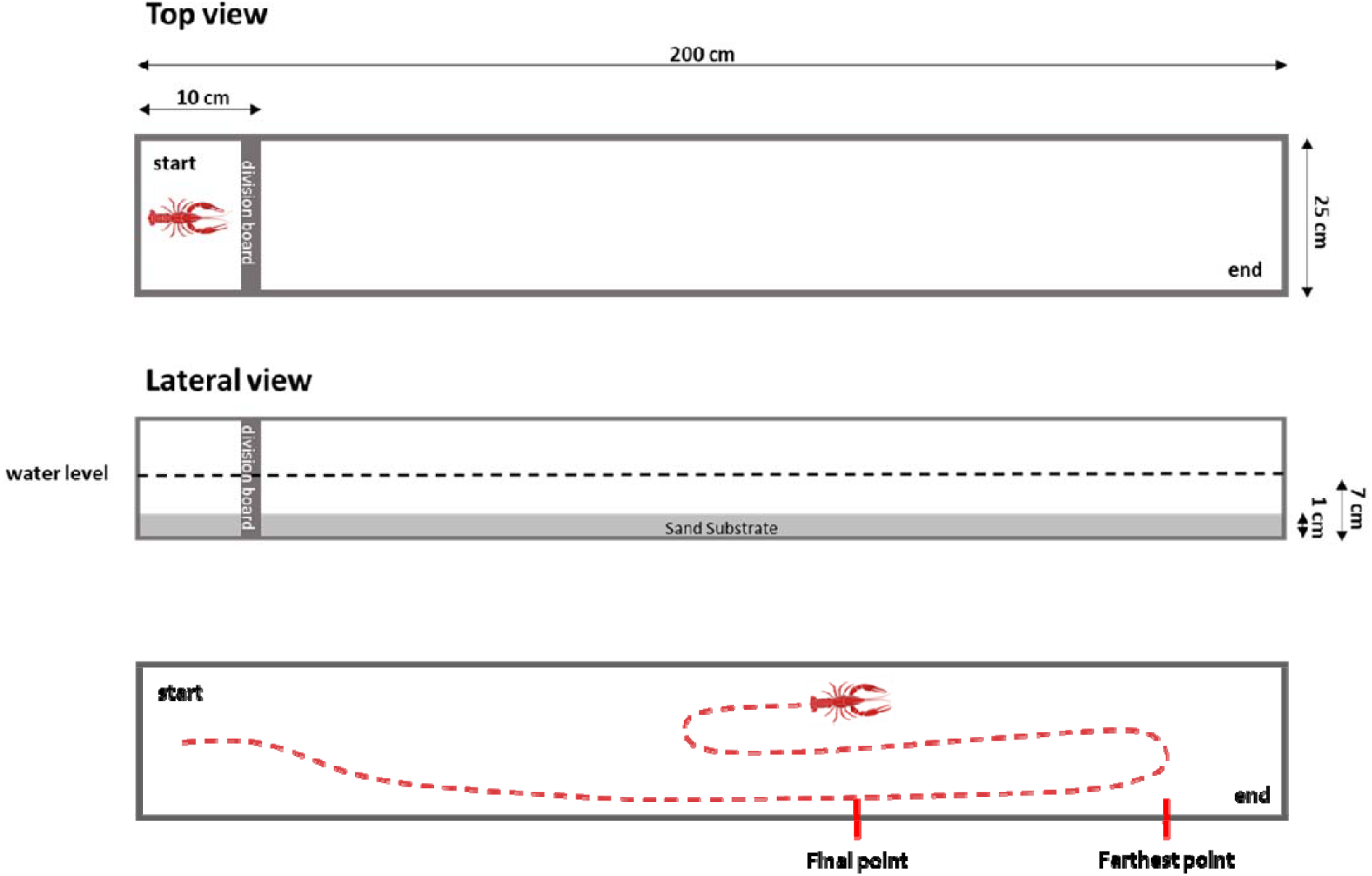
Schematic top and lateral view of the layout for the experiments. The final point corresponds to the position at which crayfish are at the end of the 2 minutes (used to calculate final point velocity). The farthest point corresponds to the farthest distance crayfish achieved during their movement (used to calculate furthest point velocity). The dotted line corresponds to the total distance travelled by the crayfish (used to calculate real velocity). The division board was a translucent plastic plate.

### Environmental factors

A total of 216 individuals were used, 106 males and 110 females, with mean cephalothorax length of 34.27 mm (sd = 12.76 mm). These were distributed evenly by 8 treatments, with 27 individuals each with an even size distribution. Unfortunately, 4 of these individuals didn’t survive, resulting in a slight imbalance on the number of crayfish per treatments. The experiment occurred over a period of 8 days. For the light treatments, we used the room’s fluorescent light bulbs. The room’s lights were turned off ensuring total darkness for the dark treatments and to visualize when a crayfish reached the end of the track, a commercial red laser pointer was used to illuminate for 1 to 2 seconds the end of the track every 15 seconds. The temperatures used were 16°C and 22°C, corresponding to common water temperatures in spring and summer, respectively, in rivers from the south of Portugal (Reis and Araujo 2016). The target temperature was set the day prior the day of experimentation. For the food treatment, a sardine cut in haft was placed at the end of the gutter 15 minutes before the start of the experiment. The foodless treatments were performed first to ensure that there were no food particles or scent that could interfere with them. Due to the dark treatments making it hard to visualise the real distances travelled by the individuals, only the velocity values calculated with the crayfish final position on the track (final point velocity) were used for this experiment.

### Pairing experiment

164 individuals organized into 82 pairs were selected from the three captures and grouped into six treatments, males of the same size, females of the same size, males and females of the same size, different size males, different size females and different size males and females. Another 30 individuals, 9 males and 21 females, were used individually as a control group. Crayfish used in the paired treatments had a mean cephalothorax length of 37.12 mm (sd = 11.84 mm). Those used in the control had a mean of 29.92 (sd = 12.88 mm). Considering the 82 pairs, only 79 yielded results as some individuals died before being tested, making a total of 158 individuals plus the 30 in the control. These treatments were all conducted under the same light levels and in the absence of food but were divided between two temperatures, 16°C and 22°C. They occurred over a period of 4 days for each capture, adding up to 12 days in total.

The procedure was the same as the previous experiment, with 60 seconds of acclimatization at the start of the gutter, followed by 120 seconds of free roam through the entire gutter. The only difference was that it was a pair of crayfish that were introduced instead of a single individual. The interactions between crayfish were recorded using a webcam placed above the gutter.

Each crayfish in a pair was placed on either the right or left side of the track as a way of keeping track of which individual it was. Similar to the other procedures, the total distance that each individual reached at the end of the 120 seconds was noted. If one of the two individuals reached the end of the track before the two minutes passed, the time it took for that crayfish to reach the end and the position of the other individual at that moment was noted.

### Statistical analysis

All statistical analyses were done using RStudio (2023.06.1 Build 524, Posit Software, PBC) with the libraries *tidyverse, car, lme4, rstatix* and *fitdistrplus*.

All three velocity values underwent a log(x+1) transformation in order to normalize the data and fulfil the homogeneity of variances assumption required for parametric statistical analysis (function leveneTest() from the *car* package). An One-Way ANOVA (function aov()) was used to test for differences in the three velocity values between the different treatments for the environmental factors procedure and the pairing experiment, followed by post-hoc Tukey tests (function TukeyHSD()) with Bonferroni correction to determine which treatments differed.

Additional ANOVAs and post-hoc Tukey tests were used to test for differences in all three velocity values between temperatures, size categories and the relative position of the individuals with their pair. Two sample t-tests (function t.test()) were used to compare final point velocities between the two temperatures and sexes for the environmental factor and all three velocity values for the pairing experiments. An additional t-test was used to test the three velocity values between the two size categories in the pairing experiment.

A Linear Model (function lm()) with the variable’s contrasts set to sum-to-zero was used to test the effect of temperature, food and light as well as the interaction of each two and all three of these variables on velocity. The effect of sex, size (cephalothorax length) and condition (Fulton’s index) were also tested.

Other Linear Models, also with the variable’s contrasts also set to sum-to-zero were used to test the effects of the pairing experiments and size categories on velocity. These LM were performed for the original data set, as well as the sub-sets for each temperature, size category and relative position of individuals.

## Results

### Environmental factors

Significant final point velocity differences were found between treatments (One-Way ANOVA: F = 4.303; p = 0.0002). Linear model (lm) (Table 1) and t-tests showed that temperature affected final point velocity (Two Sample t-test: t = -2.710; p-value = 0.007). Crayfish at 22°C scored the highest speed, with a mean of 0.947 cm/s (sd = 1.041). Individuals at 16°C had a mean speed of 0.645 cm/s (sd = 0.816). The presence of food items also affected final point velocity (Two Sample t-test: t = 2.206; p-value = 0.029). Crayfish achieved faster speed in the absence of food, with a mean of 0.952 cm/s (sd = 1.112) (Figure 2). Mean values for final point velocity in the presence of food were 0.640 cm/s (sd = 0.712) (Figure 2). Light did not seem to influence final point velocity (Table 1) (Two Sample t-test: t = -1.796; p-value = 0.074). Sex also did not have an effect on final point velocity (Two Sample t-test: t = 1.442; p-value = 0.151) and lm (Table 1).

**Table 1:** LM results for the environmental factors experiment on final point velocity. Significant p values are displayed in bold (significant at 0.05).

| Predictors | Estimates | std. Error | t value | p |
| --- | --- | --- | --- | --- |
| (Intercept) | 0.13 | 0.04 | 3.03 | **0.003** |
| Temperature | -0.03 | 0.01 | -2.75 | **0.007** |
| Food | 0.03 | 0.01 | 2.22 | **0.027** |
| Light | -0.02 | 0.01 | -1.84 | 0.068 |
| Cephalothorax Length | 0.00 | 0.00 | 3.06 | **0.003** |
| Sex | 0.02 | 0.01 | 1.59 | 0.114 |
| Fulton's Index | -96.09 | 137.02 | -0.70 | 0.484 |
| Temperature × Food | -0.02 | 0.01 | -1.31 | 0.192 |
| Temperature × Light | 0.01 | 0.01 | 0.50 | 0.619 |
| Food × Light | -0.04 | 0.01 | -3.38 | **0.001** |
| Temperature × Food × Light | 0.00 | 0.01 | 0.29 | 0.769 |
| Observations | 210 |  |  |  |
| R<sup>2</sup> / R<sup>2</sup> adjusted | 0.176 / 0.135 |  |  |  |

**Table 2:**
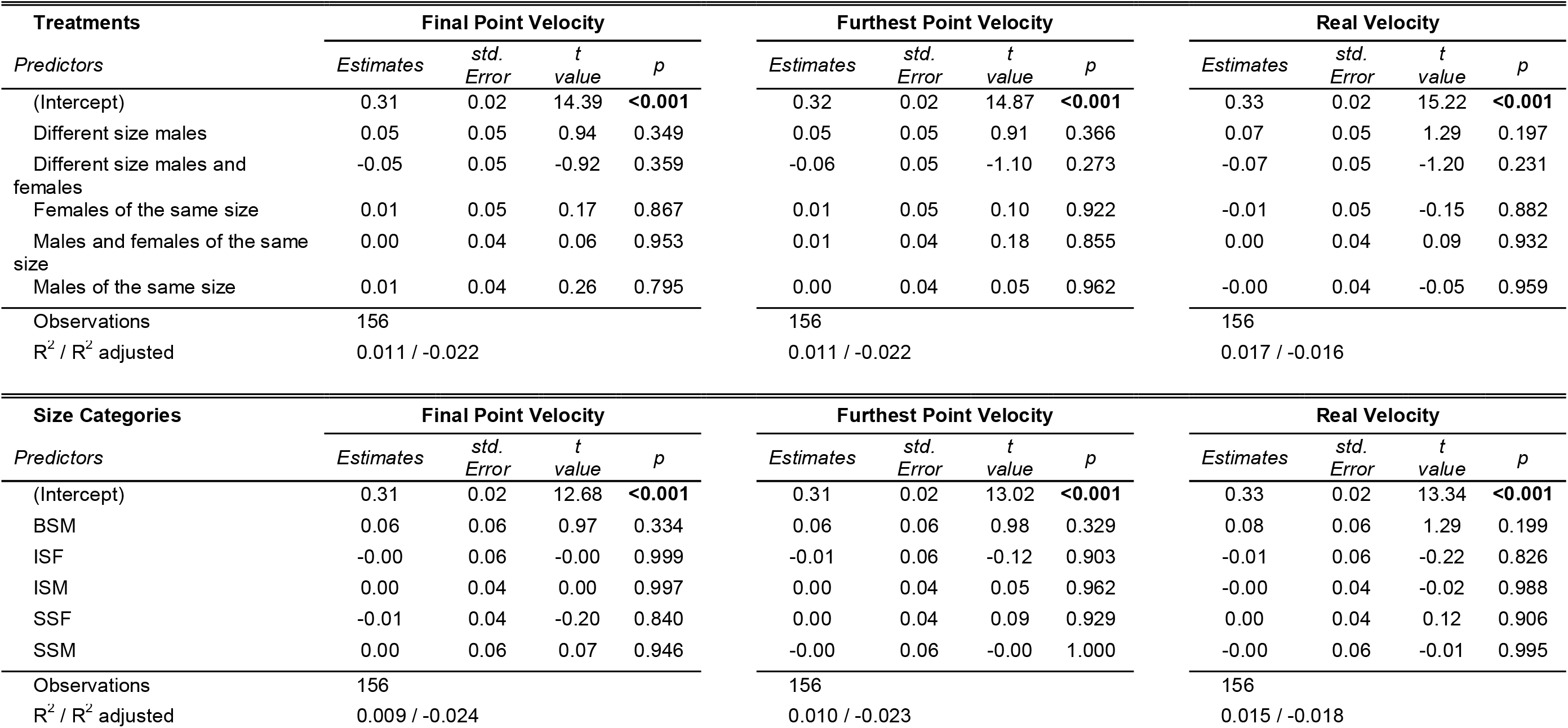
Linear Model results using the treatments and size categories from the pairing experiments to predict velocity. Final point velocity-velocity calculated using crayfish final position on the track; Furthest point velocity-velocity calculated crayfish furthest point on the track; Real velocity – velocity calculated crayfish total distance travelled.

**Figure 2:**
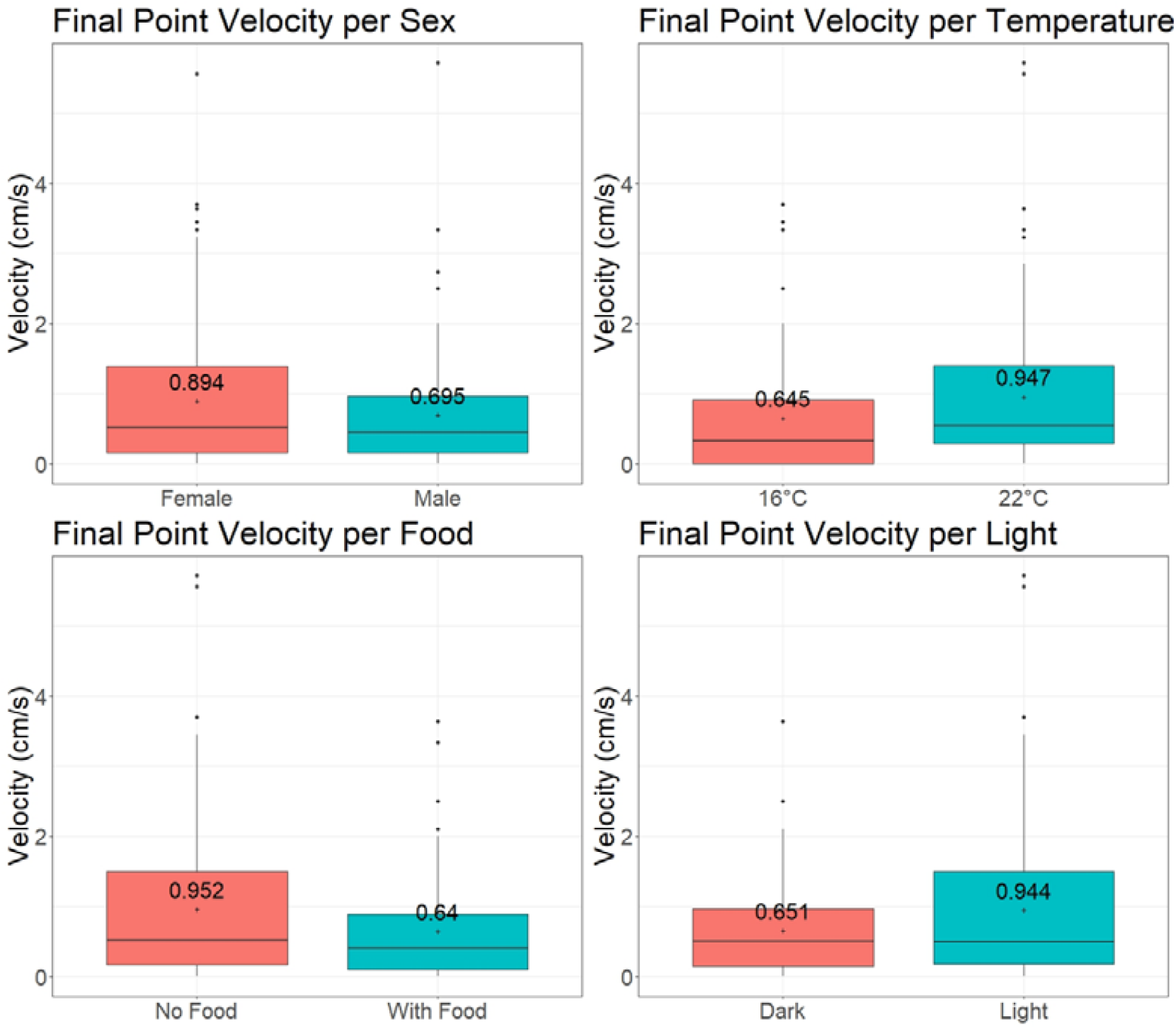
Boxplots displaying the variation of final point velocity (velocity calculated using crayfish final position on the track) from the environmental experiment between sex, temperature, food, and light. Plus sign indicates the mean value.

According to the lm, the cephalothorax length influences final point velocity (Table 1), with crayfish of [10-15[mm (mean = 0.420; Median = 0.133; sd = 0.525) and [15-20[mm (mean = 0.343; Median = 0.275; sd = 0.353) having the lowest final point velocity means, increasing up to crayfish of [25-30[mm (Mean = 1.366; Median = 0.717; sd = 1.525)(Figure 3). The velocity decreased slightly at sizes of [35-40[mm (Mean = 0.901; Median = 0.667; sd = 0.903), increased again up to sizes of [50-55[mm (Mean = 0.967; Median = 0.717; sd = 0.844)(Figure 3). There was only one individual with cephalothorax above 55 mm (Figure 3).

**Figure 3:**
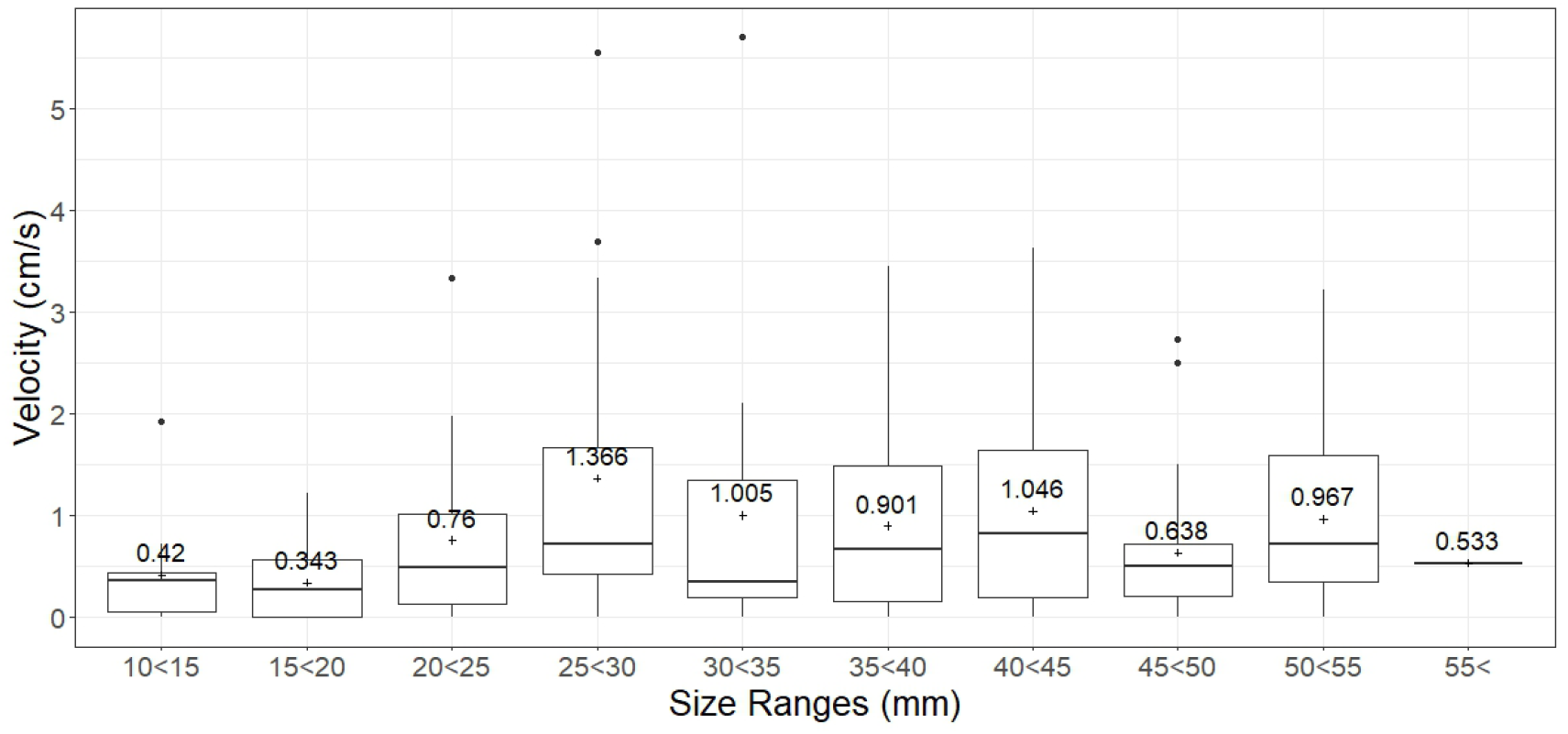
Boxplots displaying the variation of final point velocity (velocity calculated using crayfish final position on the track) from the environmental experiment testing the effect of cephalothorax length. Plus signs indicate the mean velocity.

The fastest final point velocity value was 5.714 cm/s from an individual with cephalothorax length of 30 mm in the 22°C No Food Light treatment. Both the highest mean and fastest final point velocity value were on the 22°C Without Food Light treatment (mean = 1.640 cm/s; max = 5.714; sd = 1.522). The lowest mean and max value were on the 16°C Without Food Dark treatment (mean = 0.439 cm/s; max = 1.724; sd = 0.523). Considering all individuals, the mean was 0.798 cm/s.

Also according to the lm, the interaction of temperature, food and light didn’t affect final point velocity, apart from the interaction of food with light. The light treatment with the presence of food had the lowest mean final point velocity value of 0.559 cm/s (sd = 0.657). The one with food in the dark had a mean value of 0.720 cm/s (sd = 0.76). The light treatment without food had the highest mean final point velocity value of 1.322 cm/s (sd = 1.38) and the one in the dark had a mean value of 0.582 cm/s (sd = 0.565).

### Pairing experiment

There were no significant differences in final point velocity (One-Way ANOVA: F = 0.341; p = 0.888), furthest point velocity (One-Way ANOVA: F = 0.332; p = 0.893) and real velocity (One-Way ANOVA: F = 0.506; p = 0.771). When analysing subsets for each temperature there were no statistically significant differences between treatments or size categories for any of the three velocity measurement types.

There were no significant differences between treatments at 16°C for final point velocity (One-Way ANOVA: F = 1.306; p = 0.271), furthest point velocity (One-Way ANOVA: F = 1.310; p = 0.270) and real velocity (One-Way ANOVA: F = 1.279; p = 0.282) as well as for treatments at 22°C for final point velocity (One-Way ANOVA: F = 0.558; p = 0.732), furthest point velocity (One-Way ANOVA: F = 0.642; p = 0.668) and real velocity (One-Way ANOVA: F = 0.746; p = 0.591). There was also no significant differences between size category velocities at 16°C for final point velocity (One-Way ANOVA: F = 1.353; p = 0.252), furthest point velocity (One-Way ANOVA: F = 1.326; p = 0.263) and real velocity (One-Way ANOVA: F = 1.289; p = 0.278) or at 22°C (final point velocity (One-Way ANOVA: F = 0.543; p = 0.743), furthest point velocity (One-Way ANOVA: F = 0.742; p = 0.594) and real velocity (One-Way ANOVA: F = 0.852; p = 0.517)). There were also no significant differences between sexes for final point velocity (Two Sample t-test: t = 0.658; p-value = 0.512), furthest point velocity (Two Sample t-test: t = 0.525; p-value = 0.600) and real velocity (Two Sample t-test: t = 0.626; p-value = 0.532).

Significant differences were found for final point velocity between size categories for individuals behind their pair (One-Way ANOVA: F = 2.501; p = 0.039). However, no difference was found using Tukey HSD test. There were significant differences for final point velocity between sexes among the individuals that were behind their pair inside the gutter (Two Sample t-test: t = 2.549; p-value = 0.013). Females behind their pair were on average faster than the males behind their pair (Figure 4)

**Figure 4:**
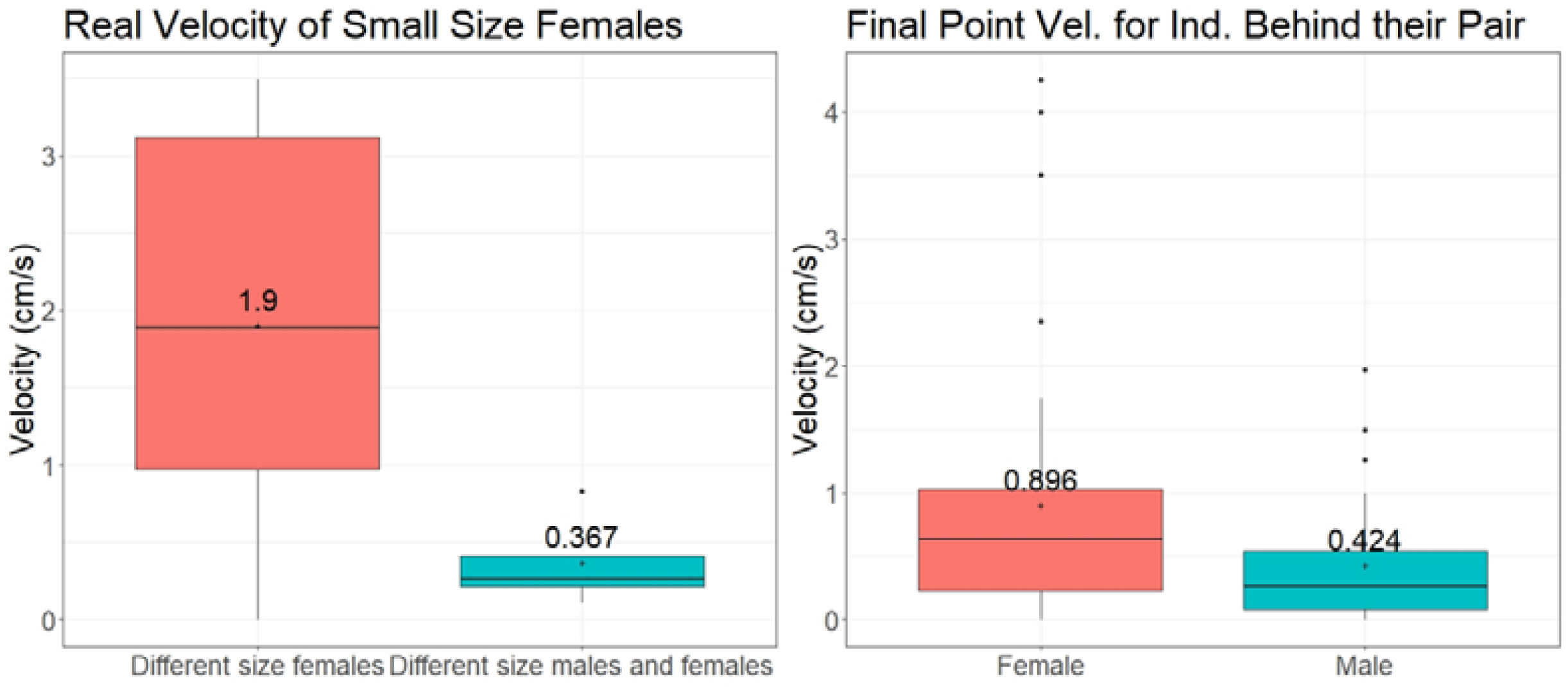
Left: Boxplot displaying the variation of real velocity (velocity calculated crayfish total distance travelled) for individuals in the different size females and different size males and females treatments in the pairing experiment; Right: Boxplot displaying the variation of final point velocity (velocity calculated using crayfish final position on the track) between males and females that were behind their pair inside the gutter in the pairing experiment. Plus sign indicates the mean value.

There were also significant differences in real velocity for small size females between the different size males and females treatments and the different size females treatment (One-Way ANOVA: F = 5.226; p = 0.0412) with the small females from the different size females treatment moving faster than those on the different size males and females treatment (Figure 4) (different size males and females had 4 individuals; different size females had 10).

## Discussion

The present paper makes a significant contribution to the understanding of the factors that influence crayfish dispersion, providing a clear perspective on the impact of temperature, food, light, and social interactions on the velocity of *P. clarkii* dispersion. As predicted, we found crayfish moving faster at 22°C than at 16°C, which lead us to suggest that with the increase in global temperatures, *P. clarkii* may increase their dispersion abilities in areas currently below the species temperature optimum.

We also predicted that food would influence velocity and that individuals would move faster in the presence of food than in the absence of it. The efficiency with which organisms detect and reach food sources is an important factor in survival, as the longer this timeframe the higher the likelihood that competitors will locate and consume the food source first. Furthermore, fresh carcasses are important sources of protein for crayfish to fuel their growth (Momot 1995). Contrary to what we expected, mean values for velocity were considerably higher in the absence of food than in its presence. The sardine carcasses used were fresh and are highly nutritive food items (Zimmer-Faust 1993) used as an effective bait to trap crayfish. All the crayfish used were starved for 48 hours before the experiments began. The attractiveness of this food item to them would be high and it was expected that individuals would be attracted to it in a laboratorial setting as they are in the wild. Crayfish can learn to have predator avoidance behaviours when presented with non-predators if these cues are paired with other alarm cues (Hazlett et al. 2002, Hazlett 2007). They can even learn to associate other species alarm cues to predators (Hazlett, 1994, 2000). It is possible that the smell of the sardine carcass, paired with the recent manipulation of the individuals as they were moved to the track, may have triggered a predator avoidance response which reduced their locomotion. If given a longer time period, we might have seen a different result as the crayfish adjusted more to the new environment of the track. Nevertheless, further research is needed on the full effects of different types of food on crayfish movement.

Because *P. clarkii* is a mostly nocturnal species (Fanjul-Moles and Prieto-Sagredo 2003), it was expected that these would move more during the dark treatments, but our results suggest that light did not affect their speed. Because this species is known to take shelter during the day to limit their exposure, the faster speeds during the light treatments might be due to individuals looking for a potential shelter or trying to reduce the time they expose themselves to potential predators that largely depend on vision for hunting (Flint 1977, Cukerzis 1988, Gherardi and Barbaresi 2000, Aquiloni et al. 2005, Marques et al. 2015).

Anastácio et al. (2015) measured the maximum speed for *P. clarkii* in nature at 255 m/12h (0.6 cm/s) and the mean at 8.8 m/day (0.01 cm/s). The same authors also measured the maximum and average speeds for *P. leniusculus*, at 461 m/12h (1.07cm/s) and 17.5 m/day (0.02 cm/s). Their results suggested that crayfish behaviour ranged from almost complete immobility, sometimes for several days, to large movements in a half day period. The speeds observed were most probably periodic bursts of locomotion as individuals look for shelter and try to avoid unnecessary exposure during light treatments.

Sex did not seem to influence the dispersal velocity according to our results, similarly to what has been reported in other studies (ex. Loureiro et al. 2015) but seems at odds with others that found differences between sexes (ex.: Barbaresi et al. 2004). This could be explained by local variation of the crayfish population, differences in the experimental designs or differences in the phase of crayfish’s gender life cycle among studies. In our case, it was expected that males would be slower than females due to their larger chelipeds. However, this was not seen, even on the bigger individuals.

Another important factor is individual size, which seems to affect speed. For example, according to Gherardi et al. (2002) velocity is positively correlated with crayfish size with speeds ranging from 1 to 11 m/day on a section of a temporary stream. The plotting of all individuals seemed to suggest that the speed varied with size, with the individuals with cephalothorax sizes in the range of 25 mm to 30 mm achieving faster speed than other size classes. Smaller individuals seem to be the slowest, potentially due to their smaller size. Interestingly, larger individuals were not that fast, possibly due to their increased weight, especially in the case of the males which develop larger chelipeds than females. Our study recorded considerably higher speeds, with a mean of 0.798 cm/s, the equivalent of 689.47 m/day, and the highest speed of 5.714 cm/s is the equivalent of 4924.8 m/day. However, it is unlikely that these speeds can be maintained for long periods of time.

Size also plays a major role in establishing dominance in crayfish (Ranta and Lindström 1992, Figler et al. 1995, Rutherford et al. 1995, Pavey and Fielder 1996), with smaller individuals retreating without fighting or fleeing from a fight (Issa et al. 1999) and larger crayfish being able to evict smaller ones from burrows (Ranta and Lindström 1992). Hence, it was expected that the smaller individuals might achieve higher speeds as they attempt to keep distance, particularly when together with larger individuals that could pose a threat to them. Surprisingly, we observed that while in the gutter, females that were behind their pair, increase their speeds (real velocity) more than males in a similar contest situation. The reason for this is unclear. When alone, sex didn’t seem to affect the velocity. We also saw that the smaller females in the ‘different size female’s treatment’ moved faster than those in the ‘different size males and females treatment’. Again, the reason why is unclear, though it could be that the small females felt less threatened by the bigger females than by bigger males. It is also important to note that the crayfish used in the experiments were kept in individual containers and didn’t socialise with other crayfish before the experiments. According to Drozdz et al. (2006) crayfish behaviour seems to depend on prior socialization. Crayfish that have been housed individually don’t show differences in social behaviours like rearing, turning, cornering or backing away from other crayfish while individuals that are housed in pairs do (Drozdz et al. 2006). Kept alone before experiments, the few moments of socialization during the experiment may have not been enough to establish a hierarchical connection between the two individuals. If the crayfish were given a longer period to socialize and establish a hierarchical structure, we might have seen significant differences in their locomotion. It is our hope that this body of work will help inform future works and improve experimental designs as to further substantiate our understanding of this invasive species.

## Conclusion

Our results show that rising temperatures may increase *P. clarkii* dispersion by increasing their locomotion while the presence of food items seems to decrease it. The combined effects of the availability of food items with temperature was also observed to influence the dispersion abilities of the crayfish. Sex did not seem to affect their speed individually, but when in pairs, females in the presence of another individual seemed to move faster than males in a similar situation, and females that were following behind another female moved faster than males that were behind females. Size also affected speed, the smallest crayfish being the slowest, these increasing in speed with size, with the fastest crayfish having an average cephalothorax length of 25 mm to 30 mm. Faced with current global changes and habitat alterations, it’s important to continuously expand our knowledge and understanding of which factors contribute to a higher rate of invasive crayfish dispersion. Knowing so may prove instrumental in formulating new plans and strategies to limit the impacts of P. clarkii.

## Acknowledgements

M.C.S. and F.B. were supported by the Foundation for Science and Technology through an individual contract (2021.01458.CEECIND/CP1668/CT0003 and CEEC/01896/2021, respectively).

